# Modelling radio-induced peroxidation of membrane lipids at ultrahigh dose-rate with pulsed beam

**DOI:** 10.1101/2025.08.08.669260

**Authors:** Rudi Labarbe, Lucian Hotoiu, Vincent Favaudon

## Abstract

1

**Background and Purpose:** FLASH radiotherapy, a technique based on delivering large doses in a single fraction at the micro/millisecond timescale, spares normal tissues from late radiation-induced toxicity, in an oxygen-dependent process, whilst keeping full anti-tumor efficiency. The original model of physical-chemical mechanisms [5] underlying the FLASH effect was modified to include a two-compartment (aqueous/lipid) system to take into account key interfacial reactions, and the pulsed nature of the beam.

**Materials and Methods:** The model predictions were tested by showing a linear correlation between experimentally measured biological outcomes reported in the literature and the final hydroperoxyl lipid [LOOH]_f_ predicted by the model for the different irradiation timing patterns and oxygen concentrations.

**Results:** The primary, carbon-centered lipid radical [L^•^] fades away in less than 5 ms, reproducing the experimental observation. The model predicts a linear correlation of [LOOH]_f_ with the inverse of the square root of the dose rate, as experimentally observed. The predicted [LOOH]_f_ correlates with the recognition ratio of mice irradiated at different dose rates and oxygen concentrations; with zebrafish embryos mean body length for different beam timing structures; with mouse skin toxicity even with dose splitting; and with the survival of mice for different doses per pulse and average dose rates.

**Conclusions:** The proposed radio-kinetic model attempts to synthesize the experimental results for different beam timing patterns. It successfully shows a correlation between the predicted [LOOH]_f_ and the experimentally observed biological outcomes following irradiation with different dose rates, beam timing structures and oxygen concentrations.

## 2 Introduction

Current radiotherapy facilities usually deliver dose-rates around 0.1 Gy.s^-1^ and most clinical protocols usually involve 2 Gy daily fractions, cumulated to reach the tolerance limit of normal tissues undergoing irradiation. Another methodology named “FLASH” has emerged 11 years ago [1]. It consists in delivering large doses in a single fraction at the micro/millisecond time scale with intra-pulse dose rates in the range 10^6^ Gy.s^-1^. The dramatic advantage of FLASH is normal tissue sparing from late radiation-induced toxicity without modification of the anti-tumor efficiency [2–4].

We earlier proposed [5] that the physical-chemical mechanisms underlying the FLASH effect would result from a competition between, on the one hand, self-recombination (R^•^ + R^•^ → nonradical products) of a primary carbon centered radical R^•^ generated by hydrogen atom abstraction from a substrate RH and, on the other hand, oxygen uptake yielding a peroxyl radical (R^•^ + O_2_ → ROO^•^) responsible for peroxidative tissue damage [2]. At low [R^•^] (hence low dose rate), the reaction of oxygen addition dominates while at high [R^•^] (hence at FLASH dose rate), the self-recombination reaction dominates and limits the production of ROO^•^ [5].

In the original model [5], the nature of the ‘generic carbon-based biological molecule RH’ was unspecified because, at that time, there were not enough experimental results to support the hypothesis of either DNA, protein or lipid damages as target. Recently, Portier et al. [6] provided experimental evidence that sparing polyunsaturated fatty acids from peroxidation is a key event in the FLASH effect. Consistently, we focus here on lipids, and the generic substrate ‘RH’ is replaced by the lipid ‘LH’ in the equations. Furthermore, as the model is based on phospholipid membranes, we include a two-compartment (aqueous/lipid) analysis according to Wardman [7] and take into account key interfacial reactions by using the mathematical framework of Babbs et al. [8]. Finally, the area under the curve (AUC) of [LOO^•^] vs. time that was used in the initial model [5] as a proxy to the biological effect underpinning the FLASH effect, is replaced here by the final hydroperoxyl lipid concentration [LOOH]_f_ at the end of the irradiation.

What is more, the model now takes into account for the first time the pulsed nature of the beam in the equations.

We assume that the radio-induced [LOOH]_f_ is linearly correlated with the cell toxicity. To test this hypothesis, the correlation of the predicted [LOOH]_f_ with different experimentally observed biological outcomes reported in the literature is presented for different beam timing patterns (continuous and pulsed beam with different periods and duty cycles) and different oxygen concentrations.

## 3 Materials and Methods

The radio-kinetic model described in [5] was modified to focus on lipid LH chemistry. Figure 1 graphically summarizes the main elements of the model. The peroxidation of the lipid LH takes place in the lipid phase. As described in [9] and as was modelled in [5] the first step consists in OH^•^-induced hydrogen atom abstraction from LH leaving the carbon-centered radical L^•^ that can evolve into the peroxyl radical LOO^•^ by oxygen capture (Figure 1). LOO^•^ acts as initiator of a radical chain mechanism leading to amplification of lipid peroxidation. This is a first order reaction in [L^•^].

**Figure 1:**
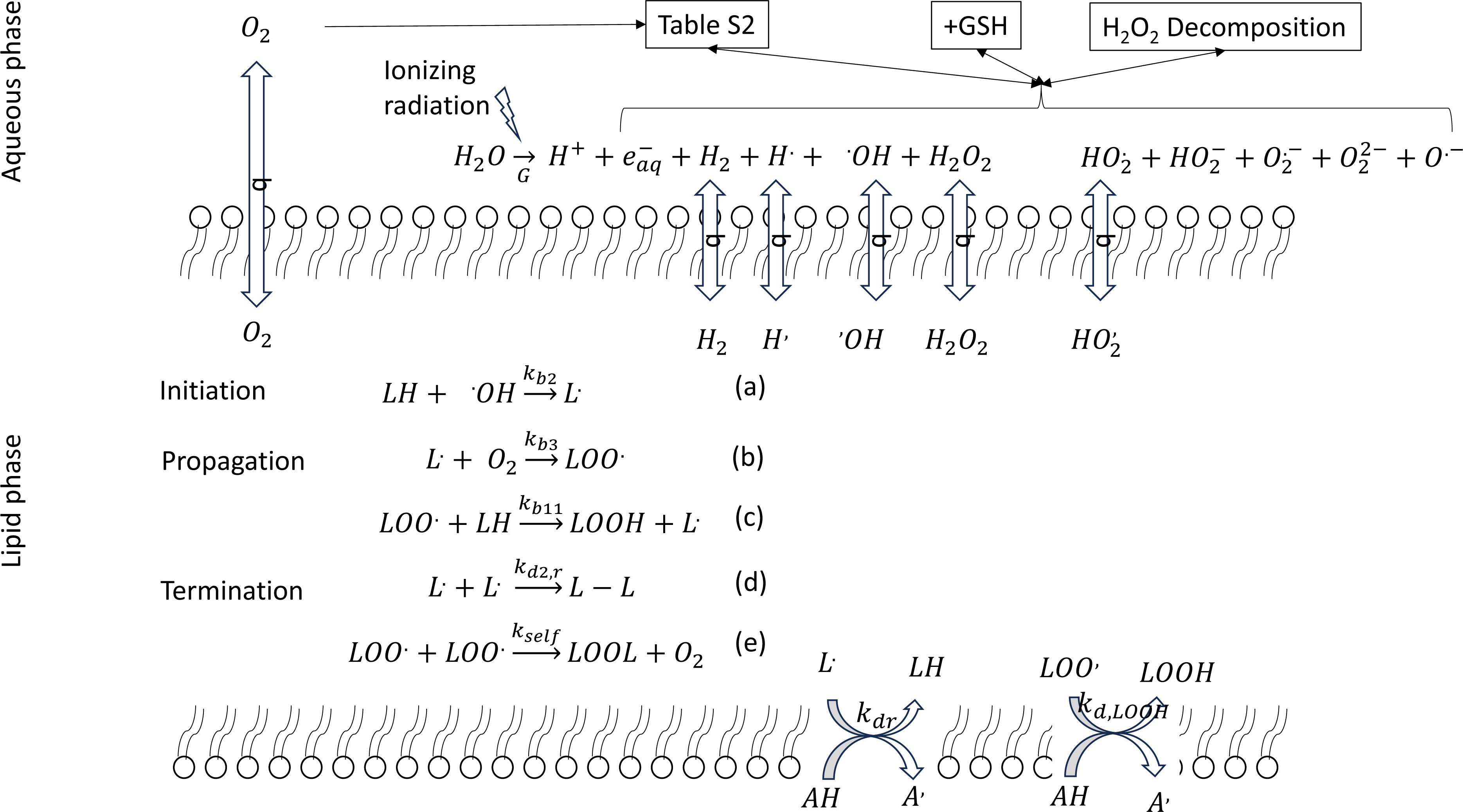
Scheme of radio-induced lipid peroxidation. The lipid phase is inside the two lipid layers schematically represented by the circles and wiggly lines. The ionizing radiation causes the decomposition of the water molecules generating radicals with radiolytic yields G specified in Supplementary Table S-1 in [5]. The radiolytic products undergo reactions with the intra-cellular oxygen in the aqueous phase leading to the creation of new species via the reactions listed in Supplementary Table S-2 in [5]. The reactions, in the aqueous phase, with thiol-based compounds (GSH) and the decomposition of H_2_O_2_ are detailed in [5]. The neutral molecules and the neutral radicals are partitioned between the aqueous and lipid phases with the partition coefficient q. In the lipid phase, the radical reactions with the lipids are grouped in the initiation, propagation and termination phases. In the termination phase, radicals L^•^ or LOO^•^ react with an antioxidant AH.

The termination takes place when two L^•^ or two LOO^•^ molecules meet and react together. Additionally, there can be termination reactions when those radicals react with an antioxidant (AH, for example, α-tocopherol, vitamin E) which leads to the formation of fatty acyl hydroperoxide LOOH [8, 10].

The rate constants used in the simulations are reported in the last column of Supplementary Table S-4.

The initial model [5] with one single phase solution was adapted to a two-phase system, (aqueous and lipid membrane), between which various chemical species are partitioned. The mathematical scheme of Babbs et al. [8] to deal with two compartments systems defines an equivalent one-phase model. The mathematical details are given in the Supplementary Materials.

As pointed out by Babbs [8], if the two reactants are confined to different compartments, the reaction rate will be zero. If both reactants are confined to a single compartment, a concentration effect occurs and the rate constant is multiplied by a factor 40. The compartmentalization of oxidizable lipids in membranes enhances the probability of chain propagation by making the effective concentrations of potential chain carriers and oxidizable lipids in the membrane much greater than their volume-averaged concentration.

In the remaining cases, the equivalent rate constant for the two-phase system is the same as for the one-phase system.

### 3.1 Simulations

The simulation assumes a beam of monoenergetic low LET radiation. The beam timing patterns are shown in Supplementary Table S-3.

The reaction rates of the ten species tracked in the model are described by a system of ordinary differential equations (ODE) (S.21 in Supplement of [5]), integrated numerically in MATLAB (R2022b), with ode15s.m using default tolerance, imposing non-negative solutions. The resolution of the ODE by ode15s.m is started at 1μs with the initial concentration of the radical species estimated from the radiolytic yields (G) of Table S-1 in Supplement of [5] and the dose delivered during the first microsecond.

To numerically solve the ODE, during each period of a beam pulse, the function ode15s.m is called twice: once for the short beam ON time (few microseconds) and a second time during the beam OFF period (several milliseconds).

The optimization of the rate constants is achieved using the constrained optimization function (*fmincon*) in MATLAB. The objective function is the sum of two terms. The first terms maximizes the value of the coefficient of determination for the linear regression between the predicted biological response (P_i_) for a set (i) of pulse timing pattern and the measured biological response (y_i,_, for example the percentage of crypt survival). The second term is a regularization term to keep the rate of oxygen depletion close to 0.45 mol/l Gy^-1^. The objective function is:

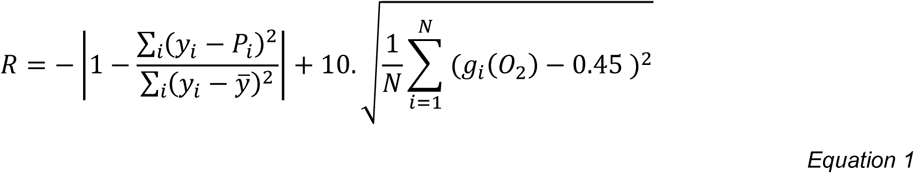

where 𝑔_𝑖_(𝑂_2_) is the predicted decrease of oxygen concentration per Gy at the end of reaction for the i^th^ set of pulse timing pattern. | | is the absolute value. The sums run over the N sets (i) of pulse timing patterns. 𝑦̅ is the average over all experimentally observed biological responses.

The source code of the program is available in open source [11].

## 4 Results

### 4.1 Rate constants

Ruan et al. [12] investigated the effect of temporal pulse structure and average dose rate of FLASH compared with CONV irradiation on acute intestinal toxicity of C3H mice with 6 MeV electrons. Figure 2 shows a plot with the predicted [LOOH]_f_ in abscissa and the experimentally measured crypt survival (Table 1 and Table 2 of [12]) in ordinate for different pulse timing patterns. The relation between [LOOH]_f_ and the observed crypt survival is unknown, but it seems reasonable to expect a sigmoid function. The sigmoid is approximated by a horizontal line at low and high [LOOH]_f_ and by a linear response for intermediate [LOOH]_f_:

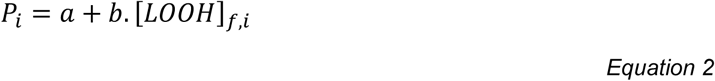

where [LOOH]_f,i_ is the final [LOOH] predicted by the model for the i^-th^ set of beam parameters and P_i_ is the predicted percentage of crypt survival. The coefficients a and b are obtained by computing in MATLAB the linear regression between the measured crypt survival (y_i_) and the predicted [LOOH]_f,i_. The linear regression line is shown in red in Figure *2*. The quality of the linear regression can be estimated by the coefficient of determination (first term of Equation 1).

**Figure 2:**
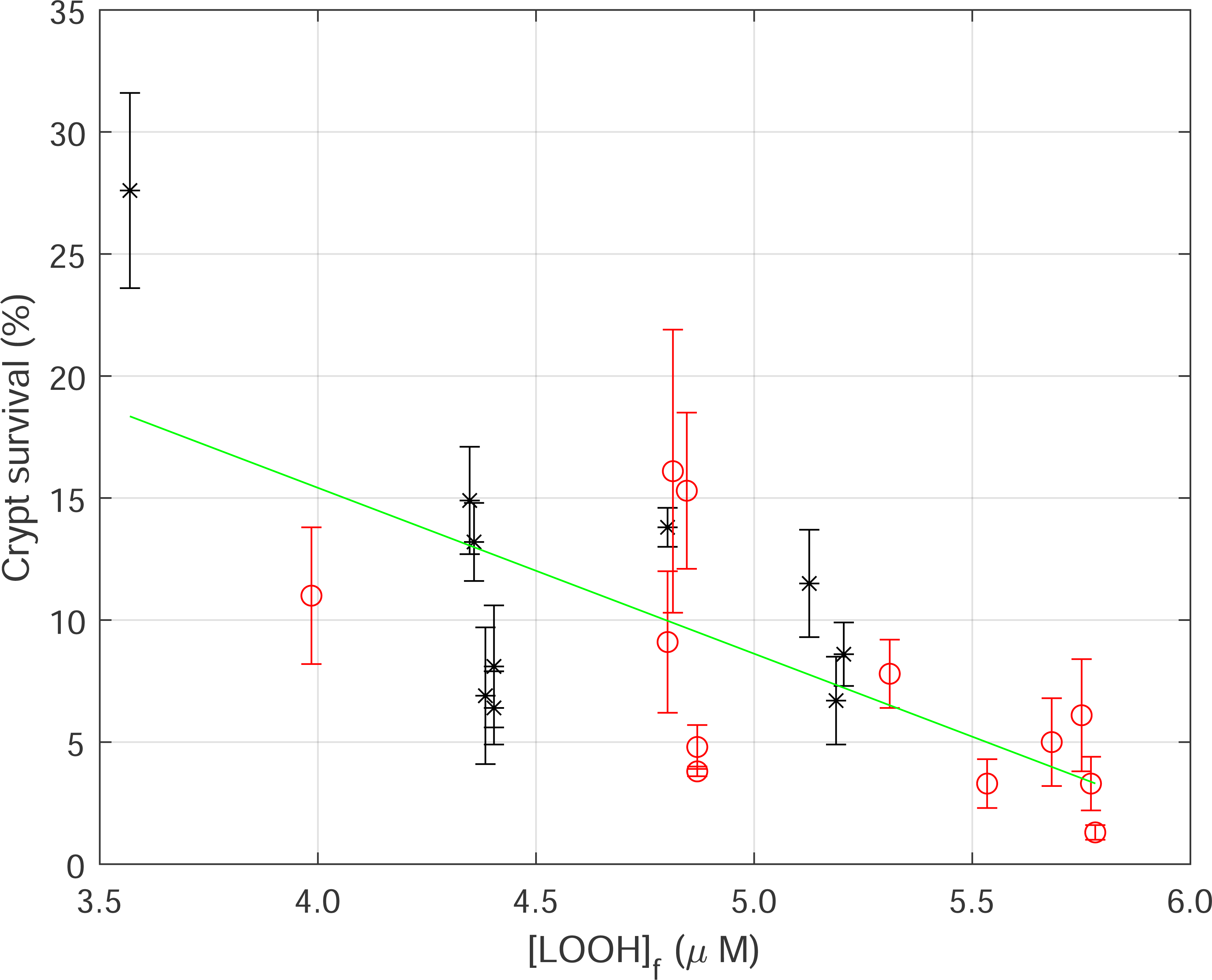
Correlation between [LOOH]_f_ predicted by the model and the percentage of intestinal crypt survival in mice exposed to whole abdominal irradiation for the beam parameters reported in Table 1 (mice aged 9 to 10 weeks, 11,2 Gy, stars) or Table 2 (mice aged 30 to 31 weeks, 12,5 Gy, circles) in [12]. The solid line represents the linear regression. Coefficient of determination: −0.4841.

The relevance of the model results depends on a sensible choice of the rate constants. A constrained non-linear optimization is carried out by varying the rate constants k_d2r_, k_b3_,k_self_, k_d,r_, k_d,LOO_. The rate constants reported in different publications (Supplementary Table S-3) are used to constrain the optimization. The optimized value of the rate constants is reported in the last column of Supplementary Table S-4.

The model predictions can now be compared to several experimental results reported in the literature.

### 4.2 Decay rate of [L^•^]

In Supplementary Figure S-5B, a 10 Gy dose, delivered with the beam parameters of the third column of Supplementary Table S-3, causes [O_2_] to decreases by < 5 μmol/L, in good agreement with Weiss et al. [13] who reported a radiation-induced oxygen consumption of 0.45 μmol/L.Gy^-1^.

The fast mixing techniques developed by Howard-Flanders and Moore [14] showed that, following irradiation at low dose rate, R^•^ fades away in absence of O_2_ with a half-reaction time of less than 5 ms in mammalian cells [15, 16]. The Supplementary Figure S-6 shows [L^•^] as a function of time predicted by the model following a single pulse (lasting 90s) with the parameters of column 1 in Supplementary Table S-3 in absence of oxygen. The Howard-Flanders and Moore [14] experiment was run at low dose rate, the [L^•^] remains low and the first order termination reaction (k_d,r_) has the main effect on the [L^•^] lifetime. The model predicts that the time after which the [L^•^] decreases below 0.2% of its initial concentration is indeed less than 5 ms.

### 4.3 [LOOH]f instead of AUC

The area under the curve (AUC) of [ROO^•^] *vs.* time was used in the initial model [5] as a proxy to the biological effect. Supplementary Figure S-7 shows final [LOOH]_f_ (Figure S-7A) and the area under the curve (AUC) of [LOO^•^] (Figure S7B) as a function of average dose rate for different duty cycles. The curves in both figures show very similar trends. The Supplementary Figure S-7C shows that the ratio of AUC to the [LOOH]_f_ for the different cases is constant with dose rate indicating that the AUC is proportional to the [LOOH]_f_. This can be understood as Equation (C) in Figure *1* shows that [LOOH]_f_ is obtained by integrating over time the [LOO^•^] (with [LOOH] constant).

Therefore [LOOH]_f_ can be used instead of AUC as a predictor of biological response.

### 4.4 Comparison to experimental results

Mihaljevic et al. [17] irradiated micelles of linoleic acid (LH) with gamma radiation (continuous beam) with dose rate up to 275 Gy/s and proposed that radiolytic yield 𝐺([𝐿𝑂𝑂𝐻]) decreases in inverse ratio to the square root of the dose rate (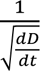).

Figure 3A is the plot of the predicted [LOOH]_f_ as a function of the inverse of the square root of the average dose rate for different pulse patterns (second column of Supplementary Table S-3) for different duty cycles. The [LOOH] _f_ was computed for different pulse periods (10 ms or 1 ms) and it is relatively independent of the pulse period. The green line shows that there is indeed a linear correlation of [LOOH]_f_ and 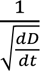 for dose rates between 10,000 Gy/s and 25 Gy/s.

**Figure 3:**
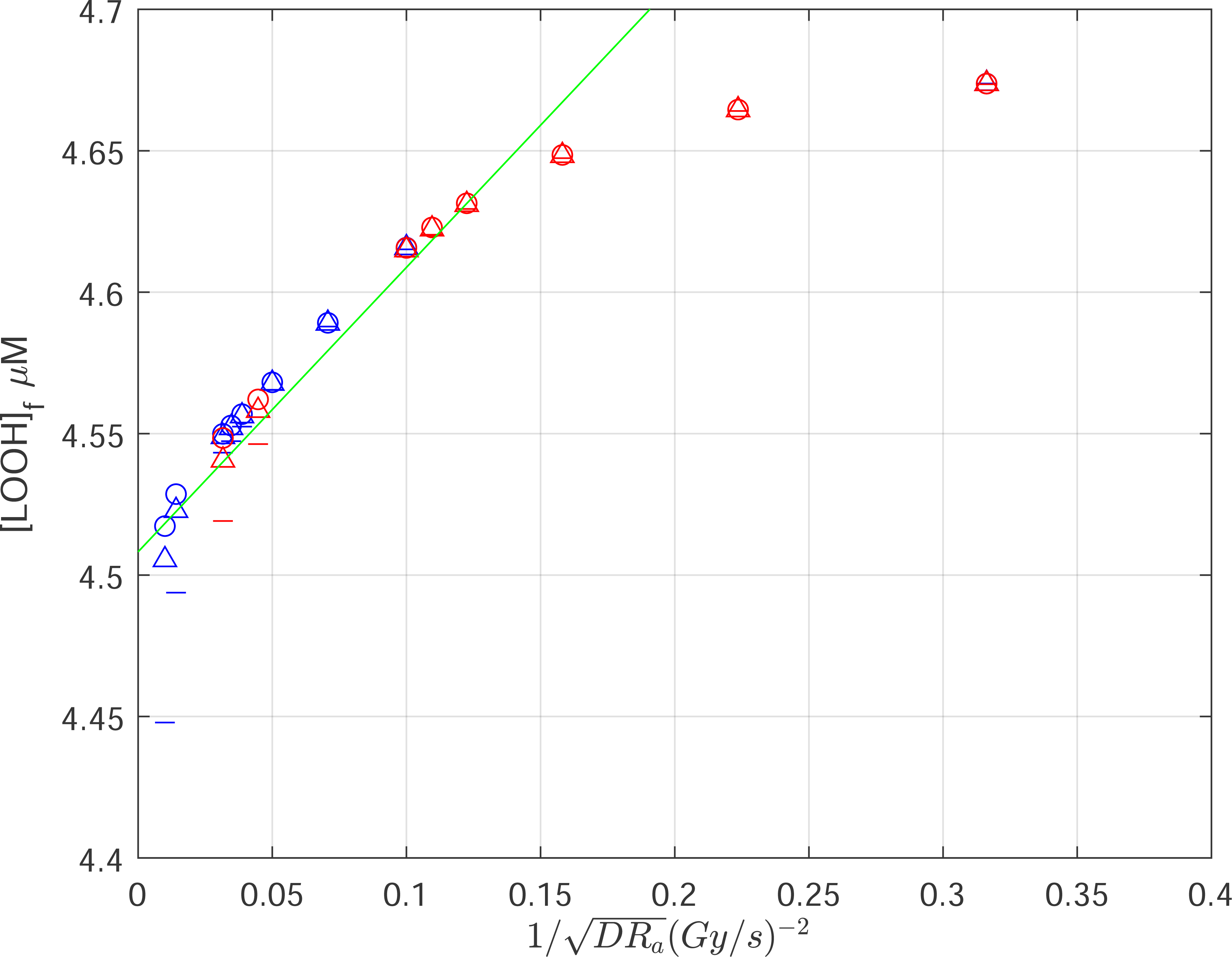
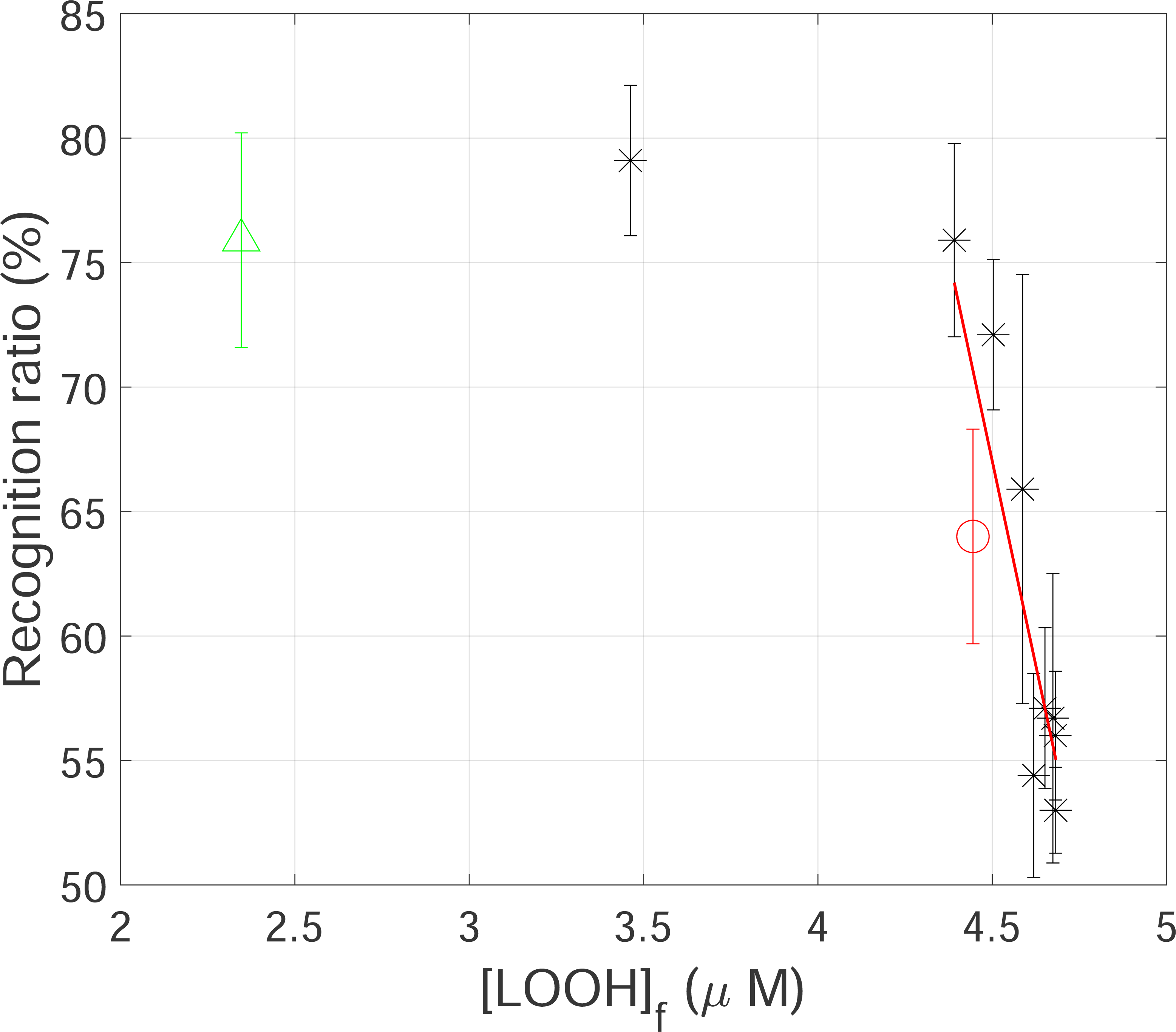
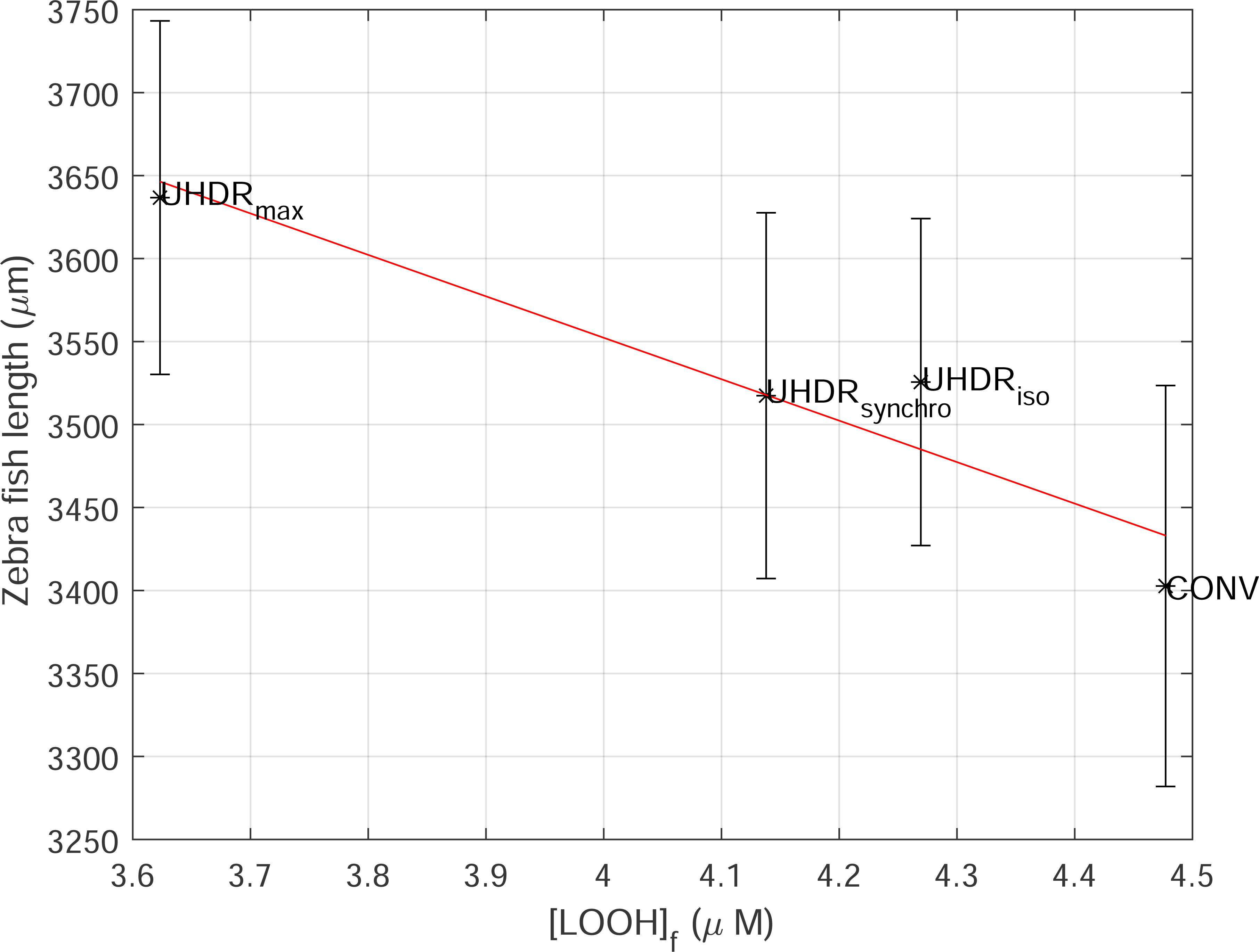
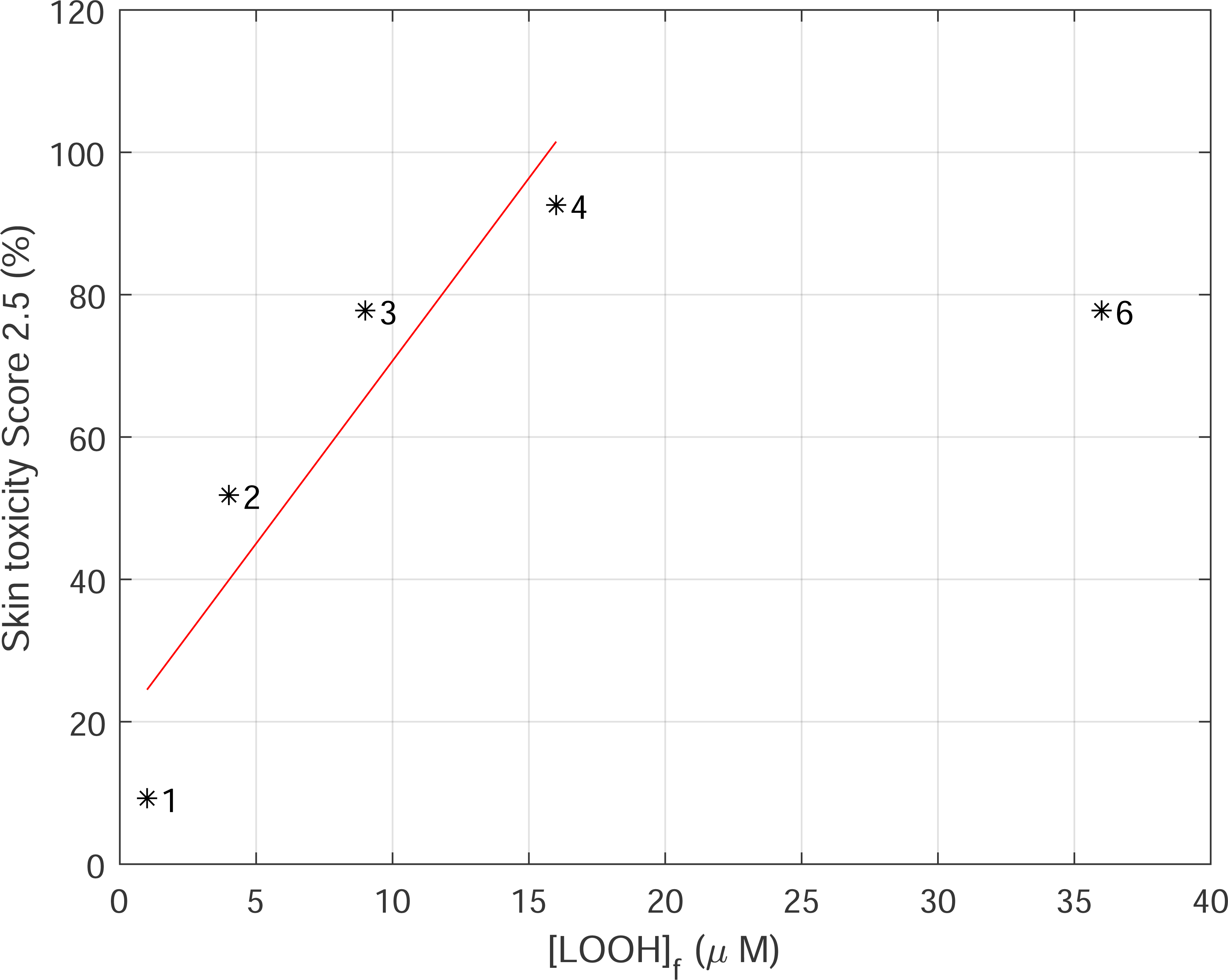
**(A)** Predicted [LOOH]_f_ as a function of the inverse square root of the average dose rate for the pulses patterns with 90% duty cycle (circle), 50% duty cycle (triangle) and 10% duty cycle (dash). The total dose is 10 Gy. The pulse period is 10 ms (red) and 1 ms (blue). The green line is a linear regression on the points in the range [0.01, 0.2 (Gy/s)^2^]. Coefficient of determination: 0.877. **(B)** Abscissa: [LOOH]_f_ predicted by the model for the beam parameters of experiment reported in figure 1C in [18] with [O_2_]_0_ = 50 μmol/l (black stars and green triangle) and in [19] with [O_2_]_0_ = 100 μmol/l (red circle). Ordinate: Measured Recognition Ratio (RR) two months post irradiation for groups of mice reported in Figure 1C of [18]. A linear regression line on the raising part of the sigmoid is shown in red. Coefficient of determination: −0.768. **(C)** Abscissa: [LOOH]_f_ predicted by the model for the beam parameters of the ELBE electron accelerator reported in Table 1 in [20]. Ordinate: Measured mean body length (mm) of Zebra fish embryo reported in Table 2 of [20]. A linear regression line is shown in red. Coefficient of determination: −0.903. **(D)** Abscissa: [LOOH]_f_ predicted by the model for the beam parameters of the proton beam reported in Table 1 in [21]. Ordinate: Measured acute damage to the skin (score 2.5) reported in figure 4A of [21]. A linear regression line is shown in red. Coefficient of determination: −0.849

Montay-Gruel et al. [18] measured the recognition ratio (RR) of mice groups receiving 10 Gy at different dose-rates, two months post-whole-brain electron-beam irradiation (pulse duration 1.8 μs, period 10 ms). In [19], they reported that doubling of the oxygen concentration in the brain during irradiation was sufficient to reverse the neurocognitive benefits of FLASH-RT. Figure *3*B shows a plot with, in abscissa, the predicted [LOOH]_f_ for the experimental conditions reported in Figure 1C in [18] with [O_2_] = 50 μmol/l (black stars) and in [19] [O_2_] = 100 μmol/l (red circle). In ordinate, the experimentally measured Recognition Ratio (RR) two months post irradiation for groups of mice as reported in Figure 1C in [18] and the RR computed from the discrimination index reported in [19] at high [O_2_]. Note that the experiment at high oxygen concentration [19] (red circle) and the single pulse experiment in [18] (green triangle) have the same pulse timing structure but different oxygen concentrations and the model correctly predicts a higher [LOOH]_f_ based on the higher oxygen concentration. A regression line is drawn through the points above [LOOH]_f_ > 4 μmol/l. Once the recognition ratio reaches 75%, the plateau of the sigmoid is reached.

Karsch et al. [20] assessed on the wild-type Zebrafish embryos mean body length, the radiobiological effects of the pulse structure of three different UHDR irradiation regimes that reflect the maximal available pulse dose rate (UHDRmax) at ELBE, the pulse structure of a clinical isochronous cyclotron (UHDRiso), a clinical synchrocyclotron (UHDRsynchro) and a quasi-continuous reference beam of conventional dose rate (CONV). The beam properties span different average and instantaneous dose rates. The beam structure reported in Table 1 of [20] was fed into the model to predict the [LOOH]_f_. Figure *3*C shows a plot with the predicted [LOOH]_f_ in abscissa and the experimentally measured embryo length (Table 2 of [20]) in ordinate showing a linear correlation between the predicted [LOOH]_f_ and the observed Zebra fish embryo length for the different pulse patterns.

Sørensen et al. [21] used a murine model of acute skin toxicity to investigate the biological effect of pauses in the dose delivered with a scanned proton beam accelerated by an isochronous cyclotron. The dose of 39.9 Gy was split into 2, 3, 4, or 6 identical deliveries with 2-minute pauses. The beam structure reported in Table 1 of [21] was fed into the model to predict the [LOOH]_f_ for each delivery of the fractional dose. The [LOOH]_f_ is computed by summing the [LOOH]_f_ of each fractional delivery. The plot (Figure *3*D) of the predicted [LOOH]_f_ vs. the experimentally measured mouse skin toxicity for different dose splits shows a linear correlation and then a plateau.

Liu et al. [22] indicated that the irradiated mice demonstrated greater survival when treated either with a high dose per pulse or with a high average dose rate irrespective of the dose per pulse. The dose per pulse was changed by pulse width modulation, while keeping the instantaneous dose rate at 1.7 10^6^ Gy/s. Figure *4*A shows the [LOOH]_f_ predicted by the model as a function of the average dose rate and of the dose per pulse. The [LOOH]_f_ depends on both parameters. For example, in Figure *4*B (resp. Figure *4*C), the [LOOH]_f_ as a function of the dose per pulse (resp. average dose rate) is plotted at constant average dose rate (resp. dose per pulse). There is a correlation between either the dose per pulse or the average dose rate and the [LOOH]_f_.

**Figure 4:**
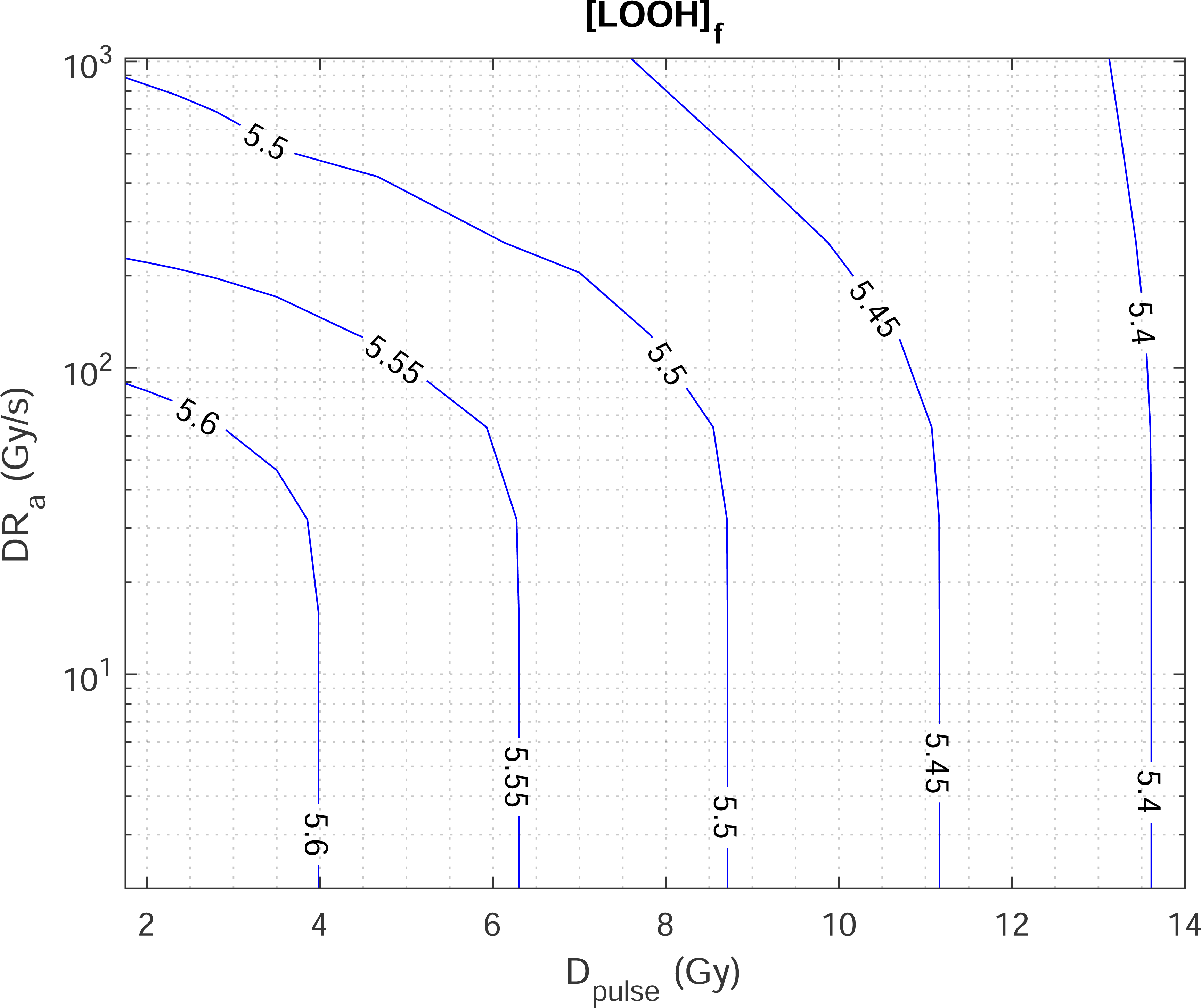
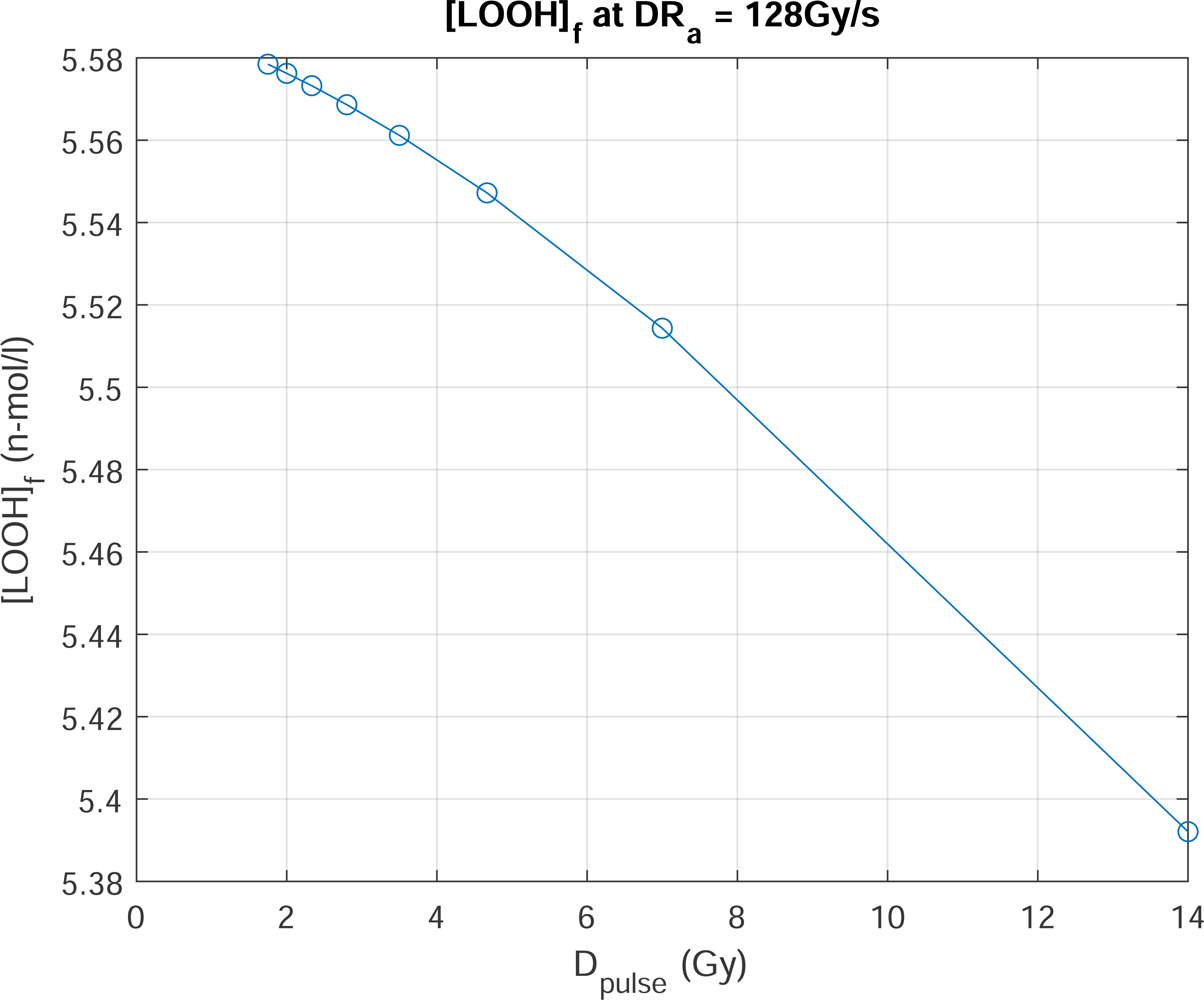
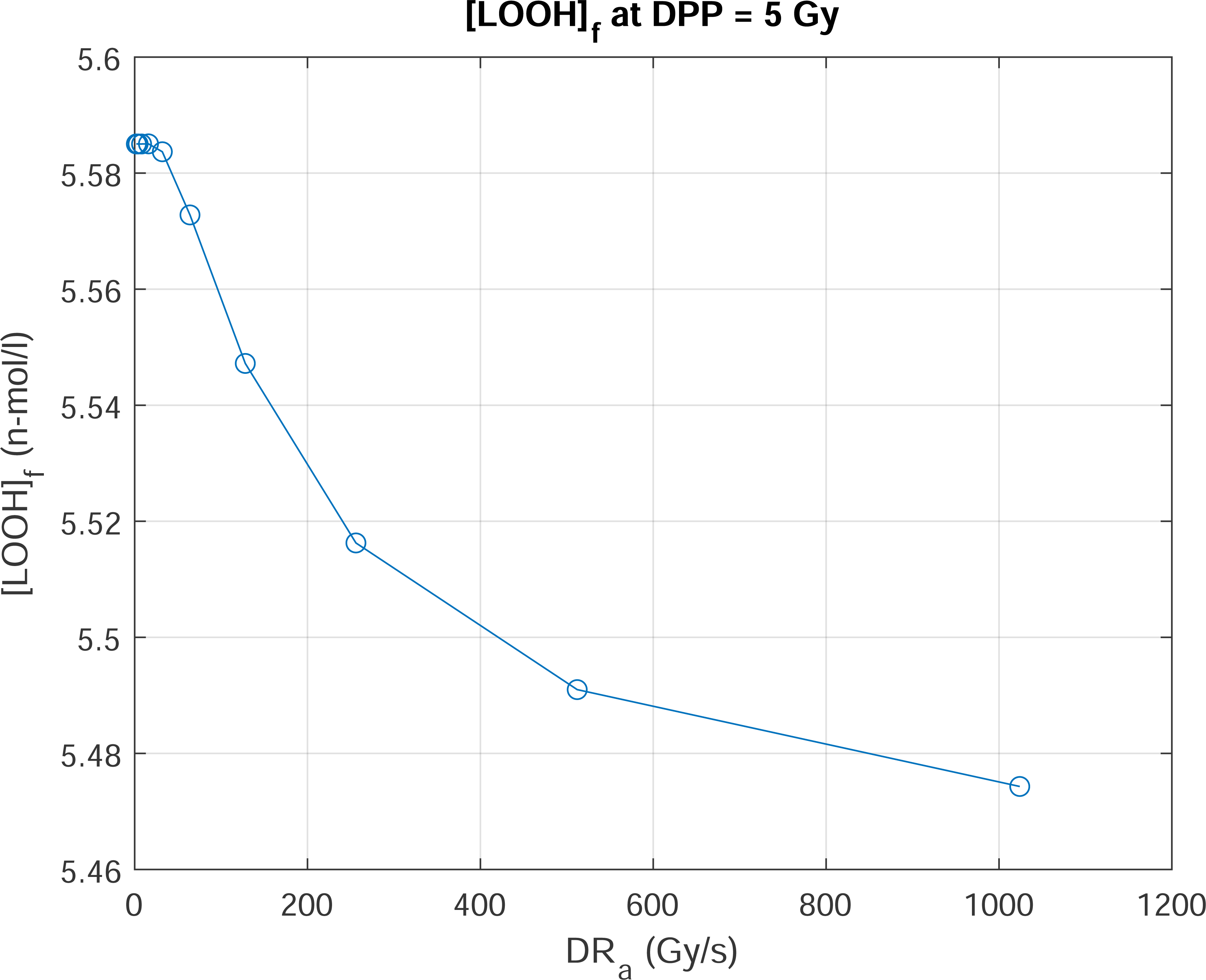
(A) Iso-concentration contour plot of predicted [LOOH]_f_ (μmol/l) as a function of the average dose rate and the dose per pulse. Total dose: 14 Gy. Peak dose rate: 1.7 10^6^ Gy/s. [O_2_] = 50 μmol/l. (B) Predicted [LOOH]_f_ as a function of the dose per pulse (at constant average dose rate of 128 Gy/s) and (C) as a function of the average dose rate at constant dose per pulse of 5 Gy. Total dose: 14 Gy. Peak dose rate 1.7 10^6^ Gy/s. [O_2_] = 50 μmol/l

## 5 Discussion

The initial kinetic model presented in [5] suggested that the FLASH effect could be explained by a competition between the first order propagation reaction of lipid peroxidation (L^•^+ O_2_) and the second order termination reaction (L^•^ + L^•^). The model predicts the final concentration of lipid hydroperoxyl [LOOH]_f_ at the end of the irradiation.

In the present paper, the original model [5] was modified to take into account, on the one hand, the pulsed structure of the beam and, on the other hand, the two-compartment description of the aqueous and lipid membrane system following the mathematical framework of Babbs et al. [8]. The value of 5 rate constants was optimized to maximize the linear correlation between the predicted [LOOH]_f_ and the experimentally measured intestinal crypt survival in mice [12]. The value of the optimized rate constants are consistent with the values reported in the literature (Table S-4).

The model predictions were successfully tested by verifying that there is a linear correlation between different experimentally measured biological outcomes and the [LOOH]_f_ predicted by the model for the different irradiation timing patterns.

Figure *2*, Figure *3*B and 3C show that there is indeed a linear correlation between the predicted [LOOH]_f_ and three different biological effects (Zebra fish embryo length, recognition ratio and crypt survival in mice) for different pulse timing patterns, (i.e. different combinations of average dose rate and peak dose rate), and for different oxygen concentrations. In addition, Figure *3*A shows that the model reproduces the observations of Mihaljevic et al. [17] that the radiolytic yield 𝐺([𝐿𝑂𝑂𝐻]) decreases in inverse ratio to the square root of the dose rate (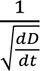) in the range of average dose rate 25 Gy/s to 10^4^ Gy/s. Supplementary Figure S-6 shows that the model reproduces the observations of Howard-Flanders and Moore [14] that, following irradiation at low dose rate, L^•^ fades away in complete absence of O_2_ with a half-reaction time of less than 5 ms in mammalian cells [15, 16].

Finally, the model predictions reproduce the observations of Liu et al. [22] that mice demonstrated greater survival when treated *<u>either</u>* with a high dose per pulse *<u>or</u>* with a high average dose rate irrespective of the dose per pulse. This suggests that the use of average dose rate or peak dose rate is probably too simplistic to be a predictor of the magnitude of the FLASH effect. A more detailed description of the beam timing pattern will be required to predict the magnitude of the FLASH effect. The radio-kinetic model proposed here may help in this attempt to synthesize the results of experimental results under different timing patterns.

## 7 Acknowledgements

This work was supported by the Walloon Region of Belgium through technology innovation partnership no. 8341 (EPT-1—Emerging Proton Therapies Phase 1) co-led by MecaTech and BioWin clusters.

VF acknowledges support from Institut Curie.

