## Supplementary material for "Modelling radio-induced peroxidation of membrane lipids at ultrahigh dose-rate with pulsed beam"

[1] IBA S.A., 3 chemin du cyclotron, B-1348 Louvain-la-Neuve, Belgium.
[2] Institut Curie, Research Division, Inserm U 1021-CNRS UMR 3347, Paris-Saclay University, PSL Research University, Centre Universitaire CS 90030, F-91598 Orsay Cedex, France.

• • •

### S-1 METHOD

A pulsed beam of monoenergetic low LET radiation leads to water radiolysis. The radiolytic yields G (mol/l/Gy) of water under anoxic conditions at 1μs post irradiation for $e_{aq}^-$, $^\bullet OH$, $H^\bullet$, $H_2$ and $H_2O_2$. were reported in Table S.1 in the Supplement of [1]

At 1μs post-irradiation, the homogeneous chemistry stage is reached. The radiolytic products undergo reactions with intra-cellular oxygen leading to the creation of new species: $HO_2^\bullet$, $O_2^{\bullet-}$ (superoxide radical), $HO_2^-$, $O_2^{2-}$ and $O^{\bullet-}$. The reaction network was described in Table S.2 in the supplement of [1]. These reactions take place in the water phase. Some of the species are involved in acid-base equilibrium as described in the Supplement S.2.1 in [1]. The aqueous phase is buffered at pH 7.0.

In addition, the decomposition of $H_2O_2$ by catalase in the water phase (equations (4) and (5) in [2]) and of $O_2^{\bullet-}$ by superoxide dismutase [1, 2] are also included.

The reactions of the hydroxyl radical $HO^\bullet$ with free radical scavengers (e.g. glutathione, GSH; reaction 8 in [1]) is also included.

As in [1] the reaction of $L^\bullet$ with free radical scavengers was also considered.

The rate constants used in the simulations are reported in the last column of Table S4.

LOOH have been studied as end-products of oxidative tissue injury [2]. The radio-induced LOOH is an indicator of the bulk peroxidation yield and damage done to the lipid membrane, which in turn elicit changes in the cell metabolism.

The initial model set in [1] with one single phase solution was adapted here to a two-phase system (aqueous and lipid membrane) between which various chemical species are partitioned with concentration $C_{aq}$ and $C_L$ in the aqueous and lipid compartment respectively. The equilibrium between the phases is assumed to be fast and is described by the partition coefficient (q):

$$q = \frac{C_L}{C_{aq}}$$

*Equation S1*

The reactants were either entirely lipid soluble ($q = \infty$ for $L^\bullet$, $LOO^\bullet$, LH ), entirely water soluble ($q = 0$ for $e_{aq}^-$, $HO_2^-$, $O^{\bullet-}$, $O_2^{\bullet-}$, $H^+$, $OH^-$), or equally soluble in both phases ($q = 1$ for $O_2$, $H_2O_2$, $^\bullet OH$, $H^\bullet$, $H_2$, $HO_2^\bullet$ ) or interact at the interphase (antioxidant).

The mathematical scheme of Babbs et al. [2] used to deal with two compartments systems, defines an equivalent one-phase model having reactant concentrations $\bar{C}$ equal to the mean, volume-averaged, reactant concentrations of the two-phase system and having reaction rate constants adjusted to account for the effects of phase separation.

To obtain the rate equations of the two-phase system (Equation S3), in the rate equations of the one phase system (Equation S2), the species concentration [A] is replaced by the mean concentration $\bar{C}_A$ and the rate constant (k) is replaced by an adjusted rate constant $k_{eq}$ :

$$-\frac{d[A]}{dt} = k.[A].[B]$$

*Equation S2*

$$-\frac{d\bar{C}_A}{dt} = k_{eq}.\bar{C}_A.\bar{C}_B$$

*Equation S3*

The total volume, $V_T$ is the sum of the volume of the aqueous compartment, $V_{aq}$, and the volume of the lipid compartment, $V_L$. The compartment volume ratio (r) is defined as:

$$r = \frac{V_L + V_{aq}}{V_L}$$

*Equation S4*

In the simulations, r is set to 40 for a typical cell [2]. The concentration of the species in each compartment can be computed from the mean concentration:

$$C_L = \frac{r}{q + r - 1}.\bar{C}$$

$$C_{aq} = \frac{q.r}{q + r - 1}.\bar{C}$$

*Equation S5*

Babbs et al. [2] assume the same rate constant, k in both the lipid and the aqueous phase. The equivalent rate constant $k_{eq}$ for the two-compartment system is (Equation 3 in [2]):

$$k_{eq} = \frac{(q_A.q_B + r - 1).r}{(q_A + r - 1).(q_A + r - 1)}.k = f(q_A, q_B, r).k$$

*Equation S6*

It is possible to solve the two-compartment model with the code of the original one compartment model [1], by transforming the compartmental concentrations to mean concentration $\bar{C}$, together with simple multiplication by the factor $f(q_A, q_B, r)$ (listed in Table S-1) of the single-phase rate constants (k) to obtain the equivalent rate constants for two-phase system ($k_{eq}$).

**Table S2**

Multiplicative coefficient $f(q_A, q_B, r)$ to convert the single-phase rate constant into the equivalent two-phase rate constant. Each line (respectively column) indicates the phase in which compound A (respectively compound B) is soluble.

| A \ B | Aqueous | Lipid | Both or interface |
|---|---|---|---|
| Aqueous | $r/_{r-1}$ = 1.02 | 0 | 1 |
| Lipid | 0 | r = 40 | 1 |
| Both or interface | 1 | 1 | 1 |

**Table S3**

Irradiation parameters used in the simulations.

| | **Single pulse, Low DR** | **Pulsed beam** | **Single pulse** |
|---|---|---|---|
| Dose | 100 Gy | 10 Gy | 10 Gy |
| Nb of pulses | 1 | From 2 to $10^4$ | 1 |
| Pulse width | 90 s | From 2 μs to 0.9 ms | 0.2 s |
| Pulse period | N.A. | 1 ms | 0.2 s |
| $[O_2]_0$ | 0 | 50 μmol/l | 50 μmol/l |

### Table S4

Comparison of the rate constants used in the different models [1] [2] [3] [4] [5]. The last but one column shows the constraints used for the optimization of the rate constants. The last column is the optimized rate constants used in the model described in this paper.

| Reaction | Rate constant | Labarbe et al. [1] | Stark et al. [3] | Babbs et al. [2] | Antunes et al. [4] | Salvador et al. Table 1 in [5] | Constraints | This model |
|---|---|---|---|---|---|---|---|---|
| **Initiation** | | | | | | | | |
| $L + {}^{\bullet}OH \rightarrow L^{\bullet}$ | $k_{b2}$ ($M^{-1}.s^{-1}$) | $10^9$ | $10^9$ | $10^9$ | $5\ 10^8$ | | | $10^9$ |
| $L + e^- \rightarrow L^{\bullet}$ | $k_{de}$ ($M^{-1}.s^{-1}$) | $1.4\ 10^8$ | | | | | | 0, different phases |
| $L + H^{\bullet} \rightarrow L^{\bullet}$ | $k_{dH}$ ($M^{-1}.s^{-1}$) | $10^8$ | | | | | | $10^8$ |
| **Propagation** | | | | | | | | |
| $L^{\bullet} + O_2 \rightarrow LOO^{\bullet}$ | $k_{b3}$ ($M^{-1}.s^{-1}$) | $5\ 10^7$ | $3\ 10^8$ | $9\ 10^6$ | $3\ 10^8$ | $2.2\ 10^8$ | [$10^6$, $3.10^8$] | $2.2\ 10^8$ |
| $LOO^{\bullet} + LH \rightarrow L^{\bullet} + LOOH$ | $k_{b11}$ ($M^{-1}.s^{-1}$) | 20 | 36 | 31 | 0.01 to 91 | $9\ 10^{-3}$ to 89.75 | | 20 |
| **Termination** | | | | | | | | |
| $L^{\bullet} + L^{\bullet} \rightarrow L\text{-}L$ | $k_{d2r}$ ($M^{-1}.s^{-1}$) | $5\ 10^7$ | $0.65\ 10^5$ | $2\ 10^8$ | | $10^6$ | [$10^5$,$10^9$] | $1.2\ 10^5$ |
| $LOO^{\bullet} + LOO^{\bullet} \rightarrow LOOL + O_2$ | $k_{self}$ ($M^{-1}.s^{-1}$) | $10^4$ | | $3\ 10^7$ | $6.6\ 10^4$ | $10^5$ | [$10^4$, $3\ 10^7$] | $1.0\ 10^4$ |
| $LOO^{\bullet} + L^{\bullet} \rightarrow LOOL$ | ($M^{-1}.s^{-1}$) | Absent | | $5\ 10^7$ | | $5\ 10^5$ | | Absent |
| Antioxidants (AH)<br>$LOO^{\bullet} + AH \rightarrow LOOH + A^{\bullet}$ | $k_{d,LOO}$ ($s^{-1}$) | k.[AH] (2) = $4.10^{-2}$ | | $10^5$<br>[AH] (2) = $50\ 10^{-6}$ M<br>k.[AH] = 5 | $5.8\ 10^3$<br>[AH] (2)=$0.2\ 10^{-3}$ M<br>k.[AH] = 1.16 | $5.8\ 10^3$ (1)<br>[AH] (2) =$0.2\ 10^{-3}$ M<br>k.[AH] = 1.16 | [0.03-10] | k.[AH] = 0.055 |
| $L^{\bullet} + AH \rightarrow A^{\bullet} + LH$ | $k_{d,r}$ ($s^{-1}$) | k.[AH] (3 = 300 | | $4.5\ 10^6$<br>[AH] (2) = $50\ 10^{-6}$ M<br>k.[AH]=225 | [AH] (3) = $1.1\ 10^{-2}$M | Absent | [30,400] | 127 |
| ${}^{\bullet}OH + AH \rightarrow A^{\bullet} + OH^-$ | $k_{dOH^{\bullet}}$ ($M^{-1}s^{-1}$) | $10^{10}$<br>[AH] (3) = $6.5\ 10^{-3}$ M | | $9\ 10^9$<br>[AH] (3) = $5\ 10^{-3}$ M | | Absent | | $10^{10}$<br>[AH] = $6.5\ 10^{-3}$ M |

---

1 Module 2 in Table 1 of [5].

2 $\alpha$-tocopherol / Vitamin E

3 Glutathione

**Figure S5**

Concentrations of (A) LOOH, LOO$^{\bullet}$, aqueous electron and (B) [$O_2$] as a function of time. The beam parameters are shown in Table 3, column 3.

**A** **B**

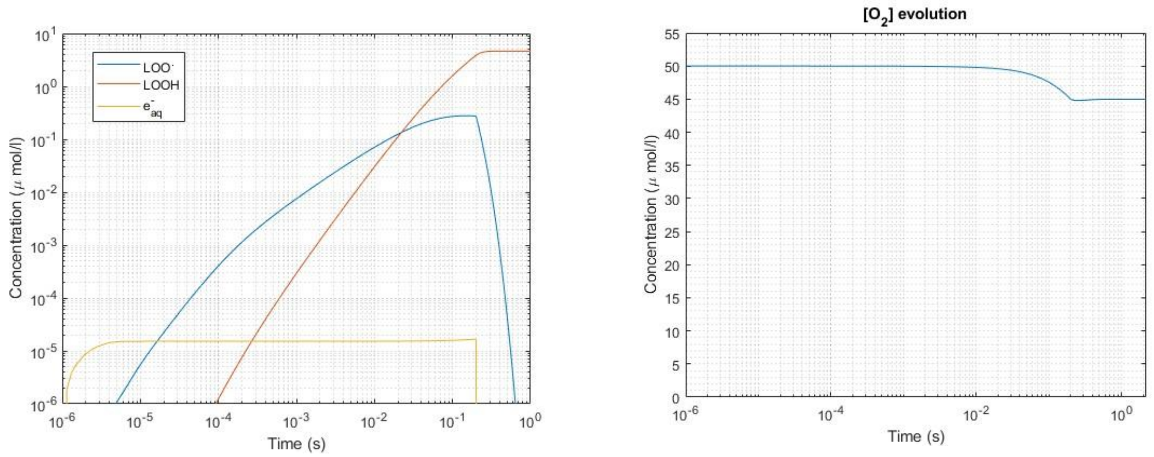


**Figure S6**

Decay of [L$^{\bullet}$] as a function of time after the end of one single pulse (lasting 90s) with the parameters of column 1 in Table S3 and using the rate constants reported in the last column of Table S3. The beam was stopped at the 90s time point, the graph represents the 5ms of the [L*] decay after the beam is stopped. In the simulations [$O_2$] = 0 mol/l.

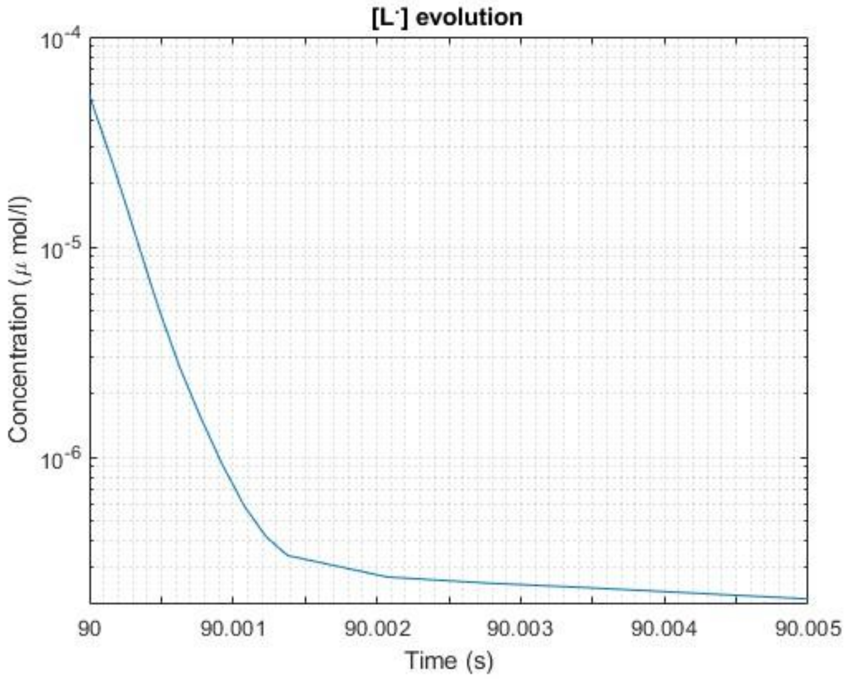

**Figure S7**

(A) $[LOOH]_f$ and (B) AUC of $[LOO^{\cdot}]$ as a function of the average dose rate (with beam parameters of second column of Table S3) and the rate constants of the last column of Table S3. (C) Ratio AUC / $[LOOH]_f$.

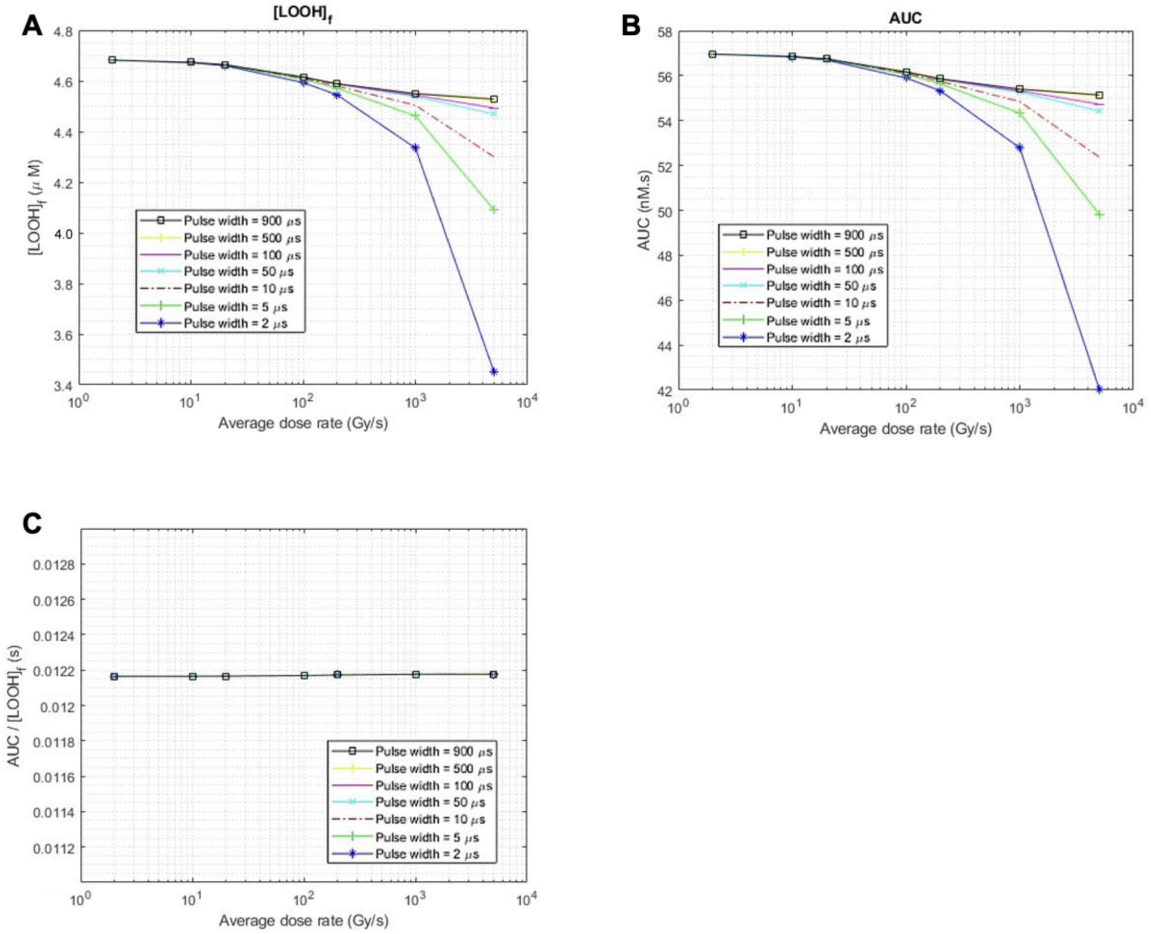

### S-8 REFERENCES


• • • • •
